# Anticipatory Gaze Reveals Individual Differences in Implicit Sequence Learning Before Motor Performance

**DOI:** 10.64898/2026.09.09.750154

**Authors:** Oindrila Sinha, Laura G Crews, William A Pleasant, Michael R Borich, Lewis A. Wheaton

## Abstract

Implicit sequence learning is commonly inferred from changes in motor performance, although anticipatory gaze behavior may provide complementary information about emerging perceptual sequence knowledge. We tracked anticipatory gaze and response time on every trial of a reach-grasp-and-move sequence-learning task to examine when individual differences became detectable in each measure.

Between-participant differences became detectable earlier in anticipatory gaze than in response time, with this ordering preserved in 95.5% of participant-level bootstrap resamples. Across the repeated sequence, anticipatory gaze became increasingly frequent, and most participants showed faster responses on anticipatory-gaze trials. Exploratory analyses further indicated that participants differed in the trial-to-trial persistence of anticipatory gaze, but these gaze characteristics were not reliably associated with sequence-specific motor learning expressed at the end of the session. Together, these findings show that anticipatory gaze and motor performance provide related but non-redundant information about implicit sequence learning where individual differences in perceptual knowledge may be detectable before corresponding differences are apparent in behavioral performance.

## Introduction

Implicit learning shapes everyday skill, and sequence learning is one of its most common forms. People improve their performance when practicing a repeating sequence of movements while remaining unable to describe, or even notice, the pattern driving the improvement^1,2^. Work in animals and humans further supports partially distinct spatial and motor representations during sequence learning, with different neural systems contributing to the acquisition and expression of sequential information^3,4^. If these components develop on different timescales, motor performance alone may provide an incomplete picture of how learning unfolds^4,5^.

Perceptual and motor aspects of a skill may develop at different rates during a training session, raising the possibility that differences between individuals may appear in one aspect before the other^6^. Evidence that perceptual and motor aspects of skill can evolve differently during learning has been reported across several tasks^6,7^. The order in which perceptual and motor changes emerge may also depend on task demands: perceptual change has been reported to precede motor change during implicit sequence learning^8^, whereas the opposite pattern has been observed during an explicit learning task^9^. These studies provide insight into the relative timing of perceptual and motor behavior at the group level, but leave an important question unresolved: when do individuals begin to differ from one another in each aspect? This distinction matters as learners who appear similar in their motor performance may already differ in the predictive information guiding their actions^10^. Quantifying when those individual differences first emerge should therefore provide a more complete account of how learning unfolds than motor performance alone.

Two obstacles have made this question difficult to test: perceptual aspects of sequence learning are rarely measured continuously at the behavioural level, and individual learners are typically collapsed into grand averages^11,12^.The first obstacle is built into the task itself. The serial reaction time task (SRTT), a standard paradigm for studying implicit sequence learning, measures motor expression through response time, but leaves the perceptual component to be inferred rather than observed^13,14^. Neural recordings have been the standard approach for probing the perceptual component, yet a recent review concludes that findings remain inconsistent across studies^15^. Anticipatory gaze offers one such behavioural measure because fixation of an upcoming target before it is cued provides trial-by-trial evidence that participants have acquired predictive information about the upcoming target location^16–18^. The second obstacle is analytic.

Group averages describe a learner who may not exist, discarding individual variation in behaviour that is structured and can be informative about later performance^11,12,19^. Here, we addressed both obstacles by recording anticipatory gaze alongside response time in a modified reach-grasp-and-move SRTT^20^ (Figure 1), providing trial-by-trial measures of anticipatory behaviour and motor expression, respectively, and by explicitly quantifying between-participant heterogeneity rather than relying only on changes in the group mean.

**Figure 1.**
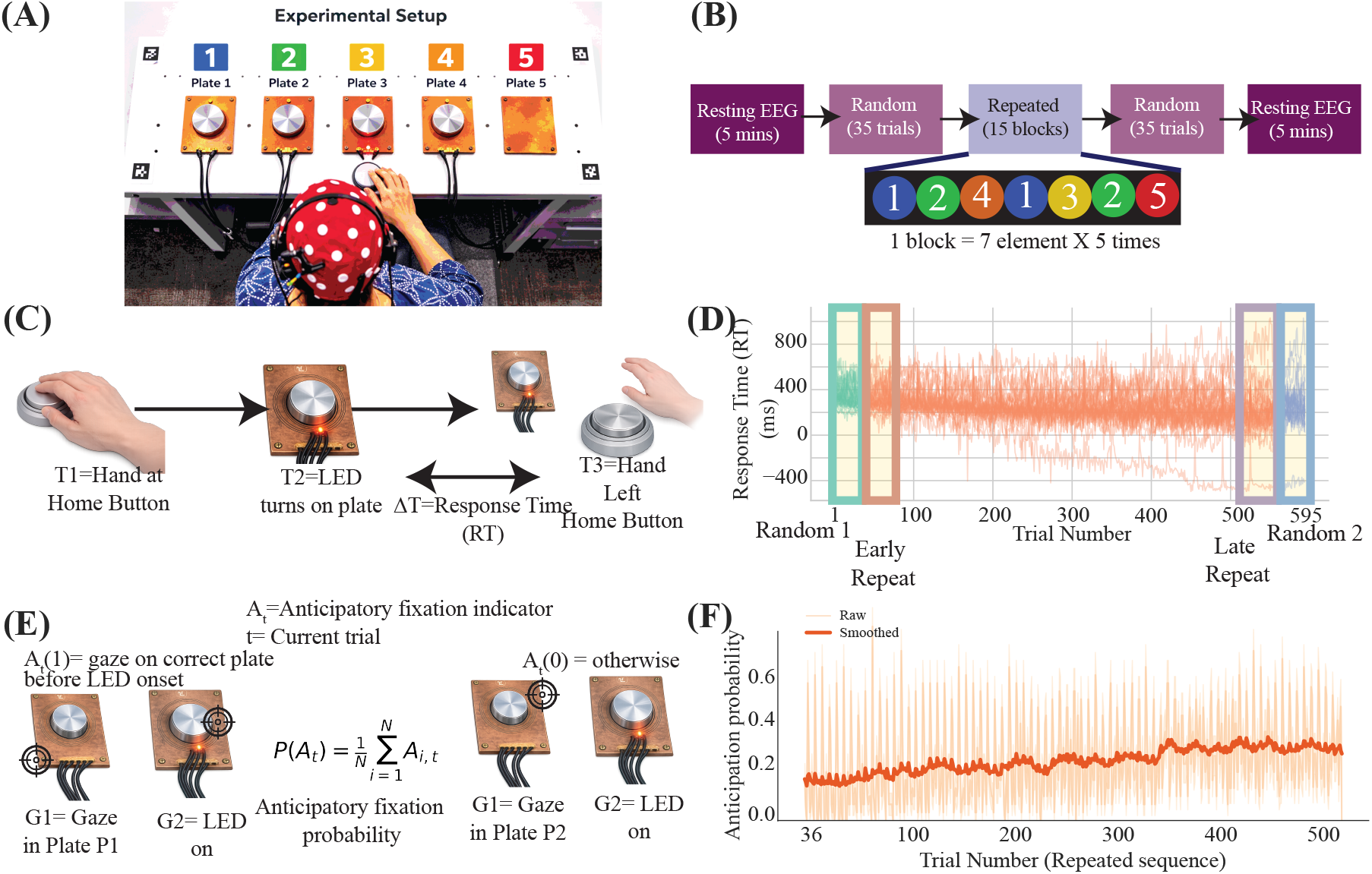
Task, paradigm and measures. (A) Apparatus: five plates in a horizontal array, four holding a disc, each with an RGB LED. (B) Session structure: resting EEG, Random 1 (35 trials), 15 repeated blocks of the 7-element sequence 1-2-4- 1-3-2-5, Random 2 (35 trials), resting EEG. (C) Trial timing: T1, hand at home button; T2, LED onset; T3, hand departure from home button. Response time RT = T3 − T2. (D) Uncleaned session-wide RT distribution across all participants, with the four analysis blocks highlighted. (E) Operationalisation of anticipatory gaze and the group anticipatory fixation probability. (F) Group anticipatory fixation probability across the repeated-sequence phase, raw and smoothed.

This approach allowed us to test two hypotheses and one exploratory question about how individual differences emerge during implicit sequence learning. First, we hypothesized that individual differences would become detectable earlier in anticipatory gaze than in response time. Second, we hypothesized that anticipatory gaze would be associated with faster responses and would become more frequent with practice. Finally, we explored whether participants differed in their coupled patterns of anticipatory gaze and motor performance across practice, forming distinct behavioural phenotypes. Together, these analyses test whether anticipatory gaze reveals individual differences before they become apparent in motor performance and characterize how those differences develop over the course of learning.

## Methods

### Participants

Thirty-five right-handed neurologically healthy adults were recruited to participate in the study. Five participants were excluded from all analyses: two had incomplete session data due to technical failures during data acquisition, and three had unusable eye-tracking data due to recording failure or failed calibration. The final analysed sample comprised 30 participants (age 24.6 ± 4.5 years, 15 M/15 F). All participants provided written informed consent prior to enrollment, and all procedures were approved by the Georgia Institute of Technology Institutional Review Board. Handedness was confirmed using the Edinburgh Handedness Inventory^21^; only participants with a laterality quotient greater than 0.6, indicating right-hand dominance, were included. Participants with three or more years of formal musical training were excluded to control for the potential influence of highly trained fine motor skills on sequence learning^22^. No participant reported a history of neurological, psychiatric, or musculoskeletal conditions affecting upper limb function.

### Experimental apparatus

Participants performed a reach-grasp-and-move modified serial reaction time task (SRTT) using five plates arranged in a straight horizontal line in front of the participant at a comfortable reaching distance (Figure 1A). Four of the five plates held a small disc; the fifth plate was left empty. Each plate contained a printed circuit board with an RGB LED to signal the target on each trial. An Arduino microcontroller (Arduino LLC) synchronised experimental events by sending simultaneous TTL pulses to a custom MATLAB program for behavioural timing, the Pupil Labs eye-tracking system, and the EEG recording system. EEG data were collected concurrently throughout the experiment but are not analysed or reported in the present paper. The reach-grasp-and-move design was selected to increase the ecological validity of the motor sequence task relative to standard button-press paradigms^20^.

### Experimental paradigm

The experiment consisted of 17 blocks of 35 trials each, yielding 595 trials in total, bracketed by five-minute resting EEG recordings (Figure 1B). At the start of each trial, the participant placed their right hand on a home button located centrally in front of the plate array. After a 500 ms delay, a red LED illuminated on one of the five plates, cueing the target. The participant then reached from the home button, grasped the disc from the illuminated plate, moved the disc to the empty plate, and returned their hand to the home button. The next trial initiated automatically once home button contact was confirmed, following a brief inter-trial interval of approximately 500 ms. No performance feedback was provided at any point during the task.

All participants completed a brief familiarisation session of five practice trials prior to the main experiment. Practice trials used a pseudorandom target sequence and were used solely to acquaint participants with the movement procedure. Following practice, participants were instructed to perform the task as quickly and accurately as possible. No mention of a repeating sequence was made at any point prior to task completion.

The first block of 35 trials used a pseudorandom target sequence with no embedded repetitions (Random 1 block) and served to establish each participant’s individual baseline performance. The subsequent 15 blocks each consisted of five complete repetitions of a fixed 7-element sequence (1-2-4-1-3-2-5; one block = 7 elements × 5 repetitions = 35 trials), yielding 525 repeated-sequence trials. The final block of 35 trials again used a pseudorandom sequence (Random 2 block), serving as the post-learning probe against which sequence-specific learning was quantified ^1,23^. A rest break of 45 seconds was provided between all blocks. The repeated 7-element sequence remained constant across all 15 training blocks and was not disclosed to participants.

Throughout, trials are numbered 1 to 525 within the repeated-sequence phase, which is the unit of all trial-resolved analyses.

### Sequence awareness assessment and participant classification

Upon completion of all 17 blocks, participants were asked two questions: first, whether they had noticed any repeating pattern during the task; and second, whether they could reproduce the sequence of movements from memory. Participants who reported noticing a repeating pattern and correctly reproduced the full 7-element sequence (1-2-4-1-3-2-5) in the correct order were classified as explicit learners^24^. Of the 30 analysed participants, five met these criteria and were classified as explicit learners. All 30 participants were retained in the primary analyses; a sensitivity analysis excluding the five explicit learners (N = 25), using the same preprocessing and analysis procedures, is reported separately.

### Behavioural data collection and trial exclusion

A custom MATLAB program recorded the timing of three events on each trial: T1, the time at which the participant’s hand was detected at the home button; T2, the onset of the target LED; and T3, the time at which the participant’s hand departed the home button to begin the reach. Response time (RT) was defined as T3 − T2 in milliseconds, the interval from cue onset to movement initiation (Figure 1C). RT therefore indexes when the movement began, not how long it took to complete. RT is used throughout in this sense.

Raw MATLAB recordings used a placeholder value of −502 to flag trials in which the timing system did not register a valid response. These sentinel values, along with any other non-finite entries, were converted to NaN and were not imputed. A physiological range filter was then applied, setting trials with RT below 50 ms or above 2000 ms to NaN. Finally, z-score outlier removal was applied within each participant, separately within Random 1, the 525-trial repeated phase, and Random 2: RT values deviating more than 2.5 standard deviations from the participant’s own mean for that block were set to NaN and excluded from all analyses. The range filter was applied before z-score removal to prevent extreme values from distorting the mean and standard deviation used for outlier detection. RTs for each block were extracted from separate named variables in the recording file rather than by position within the session, so block boundaries cannot drift. The uncleaned session-wide RT distribution, including the sentinel and out-of-range trials, is shown in Figure 1D.

Across the analysed sample (N = 30), a median of 20.0 trials per participant (3.4% of the complete 595- trial session) was excluded by this procedure, and a median of 17.0 trials (3.2%) when restricted to the 525-trial repeated-sequence phase alone. Across all remaining valid trials, RT averaged 272.8 ms (SD = 125.8 ms, range 50–1030 ms) across the complete session, and 266.9 ms (SD = 123.7 ms, range 50–1016 ms) within the repeated-sequence phase alone.

### Eye-tracking

Eye movements were recorded using a Pupil Labs wearable eye-tracking system (Pupil Labs UG, Berlin, Germany)^25^, comprising a scene camera positioned above the nasion capturing the participant’s visual field at 30 Hz and two infrared eye cameras recording pupil position from both eyes at 120 Hz, providing approximately four eye-position samples per scene frame. Gaze position within the plate workspace was estimated on each trial by mapping pupil-position data onto the scene camera image using the Pupil Labs surface tracking pipeline, which registered gaze to a calibrated region of interest corresponding to the five-plate array. Prior to the main experiment, participants fixated each plate in sequence to establish a participant-specific gaze-to-plate mapping. Calibration quality was verified by computing the spatial error between fixated and estimated gaze positions. Calibration was repeated when the spatial error exceeded 10%, and participants whose error remained above this threshold after repeated attempts were excluded from all analyses.

Anticipatory gaze was operationalised as a binary variable on each repeated-sequence trial. A trial was coded as anticipatory (Aₜ = 1) if the participant’s gaze was fixating the correct next plate at the moment of LED onset — operationally, if a fixation whose onset preceded the LED remained active at the time the LED fired (fixation start timestamp ≤ LED timestamp ≤ fixation end timestamp). A fixation already present within the correct target region at LED onset necessarily preceded the visual cue and was therefore classified as anticipatory. This operationalisation is consistent with previous sequence-learning studies in which anticipatory eye movements toward upcoming target locations provided evidence of sequence prediction^26,27^.Because classification requires a preceding inter-trial interval, the first repeated-sequence trial cannot be coded, and a small number of additional trials per participant were lost to tracking dropout.

All gaze analyses were restricted to the 525 repeated-sequence trials. Group-level anticipatory fixation probability at each trial was computed as P(Aₜ) = (1/N) Σᵢ Aᵢ,ₜ, where N is the number of participants and Aᵢ,ₜ is the binary indicator for participant i on trial t (Figure 1E–F). Across the 30 analysed participants, the group-mean anticipatory rate was 23.8% (SD = 9.5%), with individual rates ranging from 12.3% to 49.1%.

### Learning outcome measure

Sequence-specific implicit learning was quantified using the sequence-specific skill score, defined as the difference in mean RT between the Random 2 block and the final 35 trials of the repeated-sequence phase (Late Repeat). A positive score indicates that the participant was faster on the repeated sequence than on the subsequent random block, reflecting sequence-specific learning beyond general practice effects^28^. An analogous measure, the early sequence-specific skill score, defined as the difference in mean RT between Random 1 and the first 35 trials of the repeated-sequence phase (Early Repeat), was also computed and is reported alongside the sequence-specific skill score as a robustness comparison (Figure 2B). The sequence-specific skill score is used as the primary learning-outcome measure throughout, since it reflects performance after the full 525-trial training period rather than after only the first block. Block-level means for both measures were computed after applying the same three-step cleaning procedure described above. Across the 29 participants with complete block data, including the four whose scores were negative, the sequence-specific skill score had a mean of 46.9 ms (SD = 65.4 ms), a median of 33.9 ms, and ranged from −97.8 ms to 207.1 ms, with 25 of 29 (86.2%) positive. One participant had zero valid trials in the final random block after cleaning and was excluded from all skill-score-dependent analyses.

**Figure 2.**
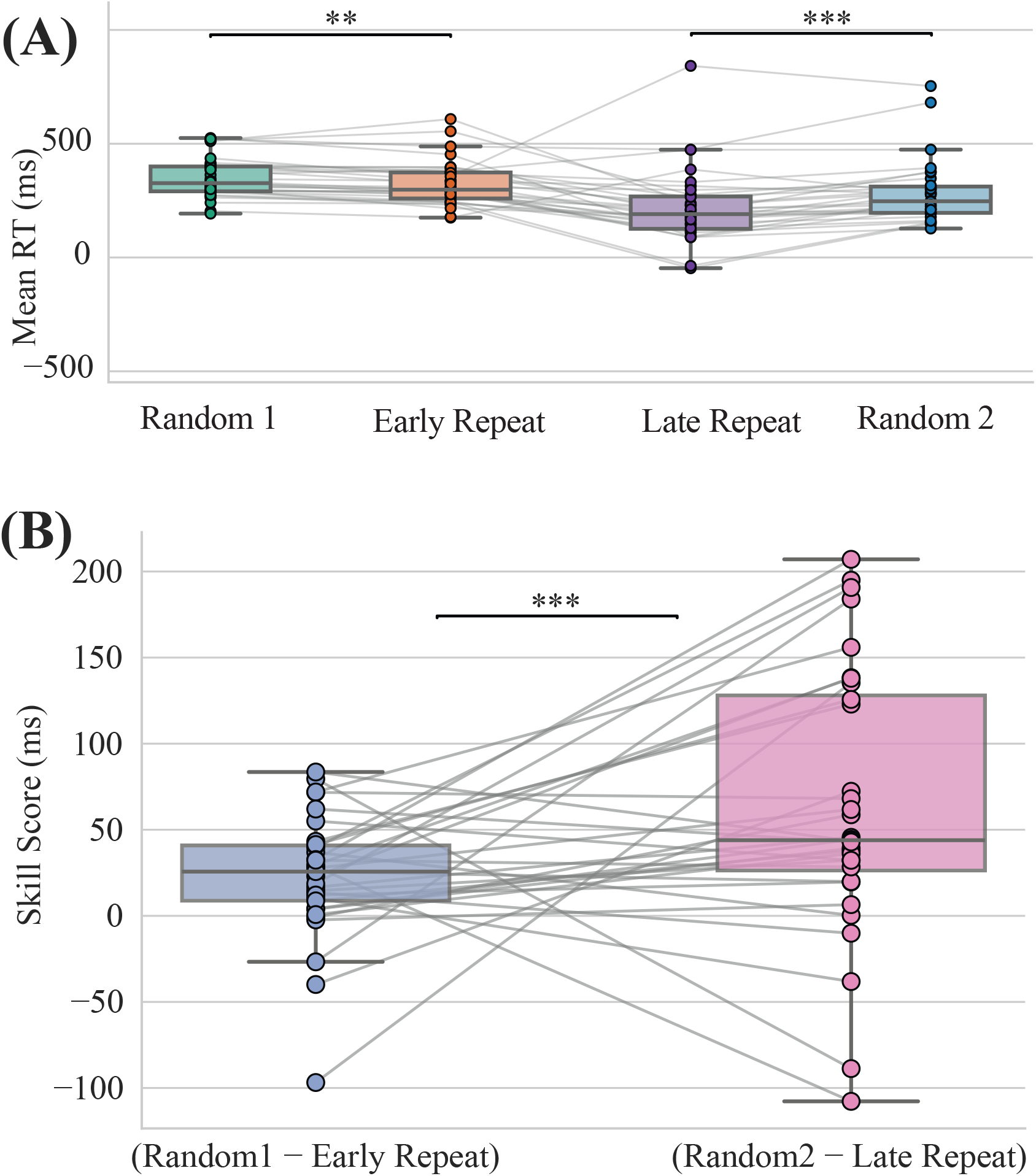
Behavioural learning. (A) Mean RT per participant across Random 1, Early Repeat, Late Repeat and Random 2, with paired-participant lines. (B) Early and late sequence-specific skill scores per participant. ** p < 0.01, *** p < 0.001.

### Statistical analysis

#### Software and analysis environment

All statistical analyses were conducted in Python (version 3.12.7) using NumPy and Pandas for data handling, SciPy (version 1.13.1) for non-parametric tests, Statsmodels (version 0.14.2) for fixed-effects regression with clustered standard errors and generalised estimating equation (GEE) models, and scikit-learn (version 1.5.1) for K-means clustering, silhouette scoring, and principal component analysis. Permutation and bootstrap procedures used NumPy random number generation with a fixed seed (42).

All analysis code is available at https://doi.org/10.17605/OSF.IO/UPD5F

#### Sample size and power

A formal a priori power analysis was not conducted. The individual-difference phenotype and clustering analyses were exploratory, and no established effect-size benchmarks are available for these analyses in the prior literature. Sample size is reported transparently alongside effect sizes and confidence intervals throughout, consistent with recommendations for sample-size justification and estimation-focused inference^29,30^.

#### Behavioural learning

Changes in response time across the session were assessed using paired-samples *t*-tests comparing Random 1 with Early Repeat, Early Repeat with Late Repeat, and Late Repeat with Random 2 among participants with complete data for all four blocks.

#### Between-participant heterogeneity and phase definition

This analysis tested whether individual differences became detectable earlier in anticipatory gaze than in response time. Heterogeneity in anticipatory gaze and RT was estimated in identical 35-trial windows advancing in steps of 7 trials. Within each window, a participant contributed data only if they had at least 20 valid gaze trials and at least 20 valid RT trials, and windows were retained only if at least 10 participants met this criterion; the same participants therefore contributed to both curves in every window.

Anticipatory gaze is binary on each trial, and for a binary variable the across-participant standard deviation is exactly √(p(1−p)), where p is the group mean. It cannot distinguish genuine between-participant heterogeneity from a change in the mean rate. Gaze heterogeneity was therefore estimated with a beta-binomial model^31^: within each window, participant i contributed kᵢ anticipatory trials from nᵢ valid trials, with latent probabilities pᵢ ∼ Beta(α, β) and kᵢ ∼ Binomial(nᵢ, pᵢ). Parameters were estimated by maximum likelihood, and heterogeneity was reported as the standard deviation of the latent probabilities, √(μ(1−μ)/(α+β+1)), which separates between-participant variation in pᵢ from binomial sampling noise within a window. RT heterogeneity was estimated as the standard deviation of participant window means after subtracting the finite-sample component: the mean of each participant’s within-window variance divided by their valid trial count was subtracted from the observed variance of participant means, and the result truncated at zero before taking the square root. Each curve was smoothed on the window grid with a Gaussian kernel (σ = 1 window) and expressed in units of its own baseline standard deviation, as (value − baseline mean) / baseline standard deviation, with the baseline defined by windows whose midpoints fell at or before Trial 35. Divergence was defined as the first window at which a curve exceeded 1.25 baseline standard deviations and remained above that threshold for three consecutive windows, corresponding to the 15-trial sustained criterion at a 7-trial step. Divergence is reported at the midpoint of the first qualifying window, so temporal resolution is limited to approximately half a window width. Throughout, divergence refers to participants diverging from one another, not to one measure separating from another. Uncertainty was quantified with 2,000 participant-level bootstrap resamples, with both curves recomputed, smoothed, standardised and re-evaluated on each resample^32^.

Resamples were classified by outcome rather than discarded: those in which both curves crossed contributed a finite difference; those in which gaze crossed but RT did not were treated as right-censored evidence that RT diverged later; the reverse case was treated symmetrically; and resamples in which neither curve crossed were recorded as undetermined.

The point-estimate divergence windows defined three empirical learning phases used in the subsequent analyses: Early (Trials 1–91), Middle (Trials 92–160), and Late (Trials 161–525). These phase boundaries were fixed before the subsequent analyses were conducted. They serve as a descriptive segmentation of the training session.

### Association between anticipatory gaze and response time

This analysis tested whether anticipatory gaze was associated with faster responses and whether the strength of this association changed with practice. The trial-level association was estimated with a fixed-effects regression fitted to all valid repeated-sequence trials:

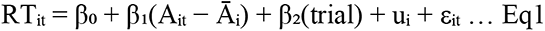

where Aᵢₜ is the binary anticipatory gaze indicator for participant i on trial t, Āᵢ is that participant’s mean anticipatory gaze rate across all 525 trials, and uᵢ is the participant fixed effect. Within-person centering removes between-participant differences in overall gaze frequency, so β₁ reflects whether a participant responded faster on trials where they looked ahead more than they typically did^33^. Standard errors were clustered by participant throughout, with a finite-cluster correction applied^34^. Equation 1 was used to characterize the gaze–RT association in three complementary ways: its consistency across participants, its average magnitude, and whether its magnitude changed across practice.

### Consistency of the gaze–RT association across participants

Each participant was reduced to a single independent estimate. For every participant, an ordinary least squares model was fitted to that participant’s own trials, RTₜ = b₀ᵢ + b₁ᵢAₜ + b₂ᵢ(trial), so that b₁ᵢ expresses their gaze-related RT difference after removing their own linear practice trend. The raw difference between that participant’s mean RT on anticipatory and non-anticipatory trials was computed alongside it as a robustness estimate. Participants contributing fewer than 10 trials in either gaze state were excluded. Because the prediction was directional, the resulting participant-level values were tested against zero with a one-sided Wilcoxon signed-rank test and an exact binomial sign test; participants, rather than trials or windows, are the independent units in these tests. Two-sided p-values were also retained for the individual participant estimates to assess whether any single participant’s coefficient differed from zero.

### Average magnitude of the gaze–RT association

To estimate the average magnitude of the association, Equation 1 was fitted jointly to all valid trials from all participants. The coefficient β₁ and its 95% confidence interval estimated the average within-participant RT difference associated with anticipatory gaze.

### Change in the gaze–RT association across practice

To determine whether the association changed with practice, a gaze × phase interaction was added to the pooled model, with Early as the reference phase. Phase-specific slopes were obtained as linear combinations of the model coefficients, and pairwise differences between phases were tested as further linear combinations. The two gaze × phase interaction terms were evaluated jointly with a Wald test as the omnibus test of change across phases. Phase-specific slopes and their 95% confidence intervals were also calculated. For visualization of the gaze–RT association across practice, Equation 1 was additionally fitted within 35-trial sliding windows advancing in steps of 7 trials. Because consecutive windows shared 28 of 35 trials, these estimates were used only to visualize the trajectory and not for statistical inference.

#### Change in anticipatory gaze frequency across practice

This analysis tested whether anticipatory gaze became more frequent as practice progressed. Separately from the gaze × phase interaction, the proportion of anticipatory trials was computed for each participant within each of the three phases.

Participants with a valid rate in all three phases were retained, so the same individuals contributed to every phase, and the relative change was expressed as the percentage increase in the group mean rate from the Early to the Late phase. Because the prediction was directional, the Early-to-Late change was tested with a one-sided Wilcoxon signed-rank test on the paired participant rates and an exact binomial sign test on the number of participants increasing.

### Phenotype identification

#### This analysis tested whether participants formed distinct phenotypes based on their joint anticipatory-gaze and motor-performance profiles across practice

Six participant-level features were extracted: mean anticipatory gaze rate in each of the three learning phases, RT improvement from the Early to the Late phase, mean RT across the final 35 repeated-sequence trials, and the sequence-specific skill score. Participants lacking an entire usable modality were excluded before feature construction; among the remainder, isolated missing values were replaced by the sample median and recorded in a missingness audit. Features were standardised to zero mean and unit variance. K-means clustering was applied with K evaluated at 2, 3 and 4, with K selected by silhouette score^35,36^. Principal component analysis was applied to the standardised feature matrix for visualisation. Because K-means cluster indices are arbitrary, clusters were labelled using a fixed descriptive rule: the cluster with the highest mean anticipatory-gaze rate across phases was labelled the higher-anticipation phenotype; among the remaining clusters, the cluster with the lowest mean sequence-specific skill score was labelled the low-learning phenotype; and the remaining cluster was labelled the typical phenotype. Because K-means maximises separation on the clustering features themselves, differences in those features across clusters are reported descriptively and cannot serve as validation. Mean RT trajectories for each phenotype were computed in sliding windows across the repeated phase and Gaussian-smoothed for visualisation.

### Gaze state transitions

#### This analysis tested whether the phenotypes differed in how anticipatory gaze was initiated and sustained across consecutive trials

For each participant, on every consecutive pair of repeated-sequence trials, the binary gaze state was recorded on trial t and trial t+1. Only genuinely consecutive pairs were used, since excluded trials break adjacency. Two transition probabilities were derived: persistence, P(anticipatory at t+1 | anticipatory at t), and initiation, P(anticipatory at t+1 | non-anticipatory at t). Both were computed in sliding windows (40 trials, step 7) and Gaussian-smoothed for visualisation. A GEE logistic model with an exchangeable working correlation structure^37^ tested the phenotype × current-gaze- state interaction, with next-trial gaze state as the binary outcome, typical phenotype as the reference phenotype, and current gaze state, phenotype, trial number and their interactions as fixed effects. Neither transition probability contributed to cluster formation, so this comparison is independent of the features defining the groups.

### Relationship between phenotype and learning outcome

#### This analysis tested whether the identified phenotypes, or the gaze-transition characteristics that distinguished them, were associated with sequence-specific learning

Because the sequence-specific skill score was itself one of the six clustering features, comparisons of skill score across the six-feature phenotypes are descriptive rather than predictive and are reported as such.

To characterize the magnitude and uncertainty of the skill-score separation within the resulting phenotype solution, we compared the higher-anticipation and typical phenotypes descriptively using complementary permutation and Bayesian summaries. The observed between-phenotype difference in mean skill score was compared with a two-sided null distribution generated by randomly permuting phenotype labels 20,000 times while preserving group sizes. In parallel, the mean difference was summarized with a normal Bayesian model using a weakly informative Normal (0, 150² ms²) prior and the observed sampling variance of the two group means. The resulting posterior distribution was used to characterize the estimated difference and the posterior probability that skill score was greater in the higher-anticipation phenotype. Because skill score contributed to the clustering solution, neither analysis was interpreted as an independent test of an association between phenotype membership and learning.

Because persistence of anticipatory gaze was the transition measure that most strongly distinguished phenotypes, and because persistence never entered the clustering, participant-level persistence and initiation were correlated directly with the skill score using Spearman correlations with permutation-based p-values and Holm correction across the three planned correlations. Persistence computed over the Early phase alone was additionally regressed on the skill score while controlling for Early-phase gaze rate, with HC3 robust standard errors and a Freedman-Lane permutation test^38^ (20,000 permutations).

## Results

### Behavioural learning

Most participants showed evidence of sequence-specific learning across the session. Sequence-specific learning was quantified using a skill score, defined as the difference in response time between the final random block and the final 35 trials of the repeated sequence (Random 2 − Late Repeat, Figure 1D). A positive score therefore indicates faster responding to the learned sequence than to the subsequently introduced random sequence. Of the 29 participants with complete data for this comparison, 25 showed a positive skill score.

Response time decreased progressively across the repeated-sequence blocks and increased again when the sequence was removed (Figure 2A). Mean RT was 339.5 ms (SD = 84.6 ms) in Random 1, decreased to 305.0 ms (SD = 100.1 ms) in Early Repeat and to 251.6 ms (SD = 149.1 ms) in Late Repeat, and then increased to 297.9 ms (SD = 130.0 ms) in Random 2 (Random 1 vs. Early Repeat: *t* = 3.74, *p* = 0.0008; Early Repeat vs. Late Repeat: *t* = 2.94, *p* = 0.0065; Late Repeat vs. Random 2: *t* = −3.86, *p* = 0.0006; *n* = 29). The increase in RT when the repeating sequence was removed indicates that at least part of the improvement was specific to the sequence rather than reflecting general changes in motor performance across the session.

Skill scores varied considerably across participants, with a mean of 46.9 ms (SD = 65.4 ms), a median of 33.9 ms, and a range of −97.8 to 207.1 ms (Figure 2B). Four participants had negative scores and therefore showed no sequence-specific benefit by this measure. Thus, although sequence learning was evident at the group level, its behavioural expression differed substantially across individuals. We next examined when these individual differences first became detectable and whether they emerged earlier in anticipatory gaze than in response time.

### Individual differences in anticipatory gaze emerge before individual differences in response time

We first tested whether individual differences became detectable earlier in anticipatory gaze than in response time. To identify when participants began to differ from one another, we tracked between-participant variability in both measures across the repeated-sequence trials and identified the first sustained increase above early-session levels. Anticipatory gaze reached this divergence point at Trial 91, whereas response time did not reach it until Trial 160 (Figure 3A). Thus, between these points, participants had begun to differ in how often they anticipated the upcoming target while remaining comparatively similar in their motor response times.

**Figure 3.**
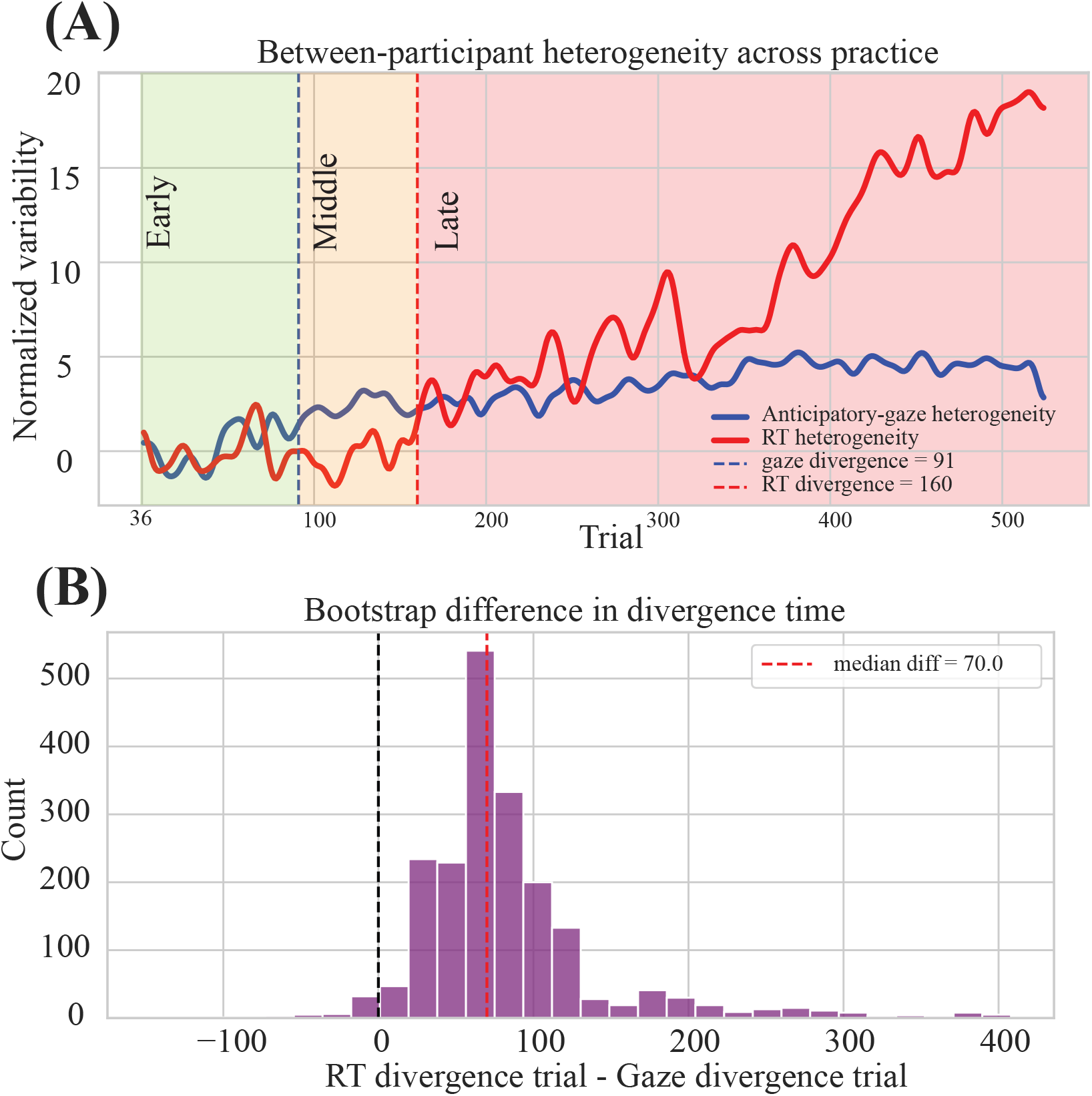
Between-participant heterogeneity in anticipatory gaze and response time. (A) Heterogeneity in each measure across the repeated-sequence phase, expressed in units of its own baseline standard deviation, with phase shading and divergence points. (B) Participant-level bootstrap distribution of the gap between the two crossings.

This ordering was preserved across participant-level bootstrap resamples. Gaze diverged before response time in 95.5% of 2,000 resamples [P(gaze earlier) = 0.955], with a median separation of 70 trials (95% CI: −15 to 330; Figure 3B). The bootstrap therefore strongly favored earlier differentiation in anticipatory gaze, although the wide interval indicates substantial uncertainty in the size of the temporal separation.

We therefore do not interpret the 70-trial difference as a precise estimate of the timing between the two transitions.

These data-derived divergence points defined three descriptive phases of the repeated-sequence period: Early (Trials 1–91), before either measure had diverged; Middle (Trials 92–160), when participants differed in anticipatory gaze but not yet in response time; and Late (Trials 161–525), when participants differed in both measures.

### Anticipatory gaze becomes more frequent across trials while its association with response time shows no detectable change

We next examined how anticipatory gaze was related to motor performance as participants progressed through the repeated sequence. Specifically, we distinguished whether the gaze–RT association changed across trials from whether anticipatory gaze itself became more frequent. Anticipatory-gaze trials were associated with faster responses in most participants, indicating a consistent direction of the within- participant association. After accounting for each participant’s change in RT across trials, 25 of 30 participants showed faster responses on trials with anticipatory gaze (median = −5.3 ms, IQR −12.4 to −0.9 ms; sign test p = 0.0002; Wilcoxon p < 0.001; Figure 4B). Because each participant contributed one estimate to these tests, this result reflects the consistency of the association across participants rather than treating individual trials as independent observations.

**Figure 4.**
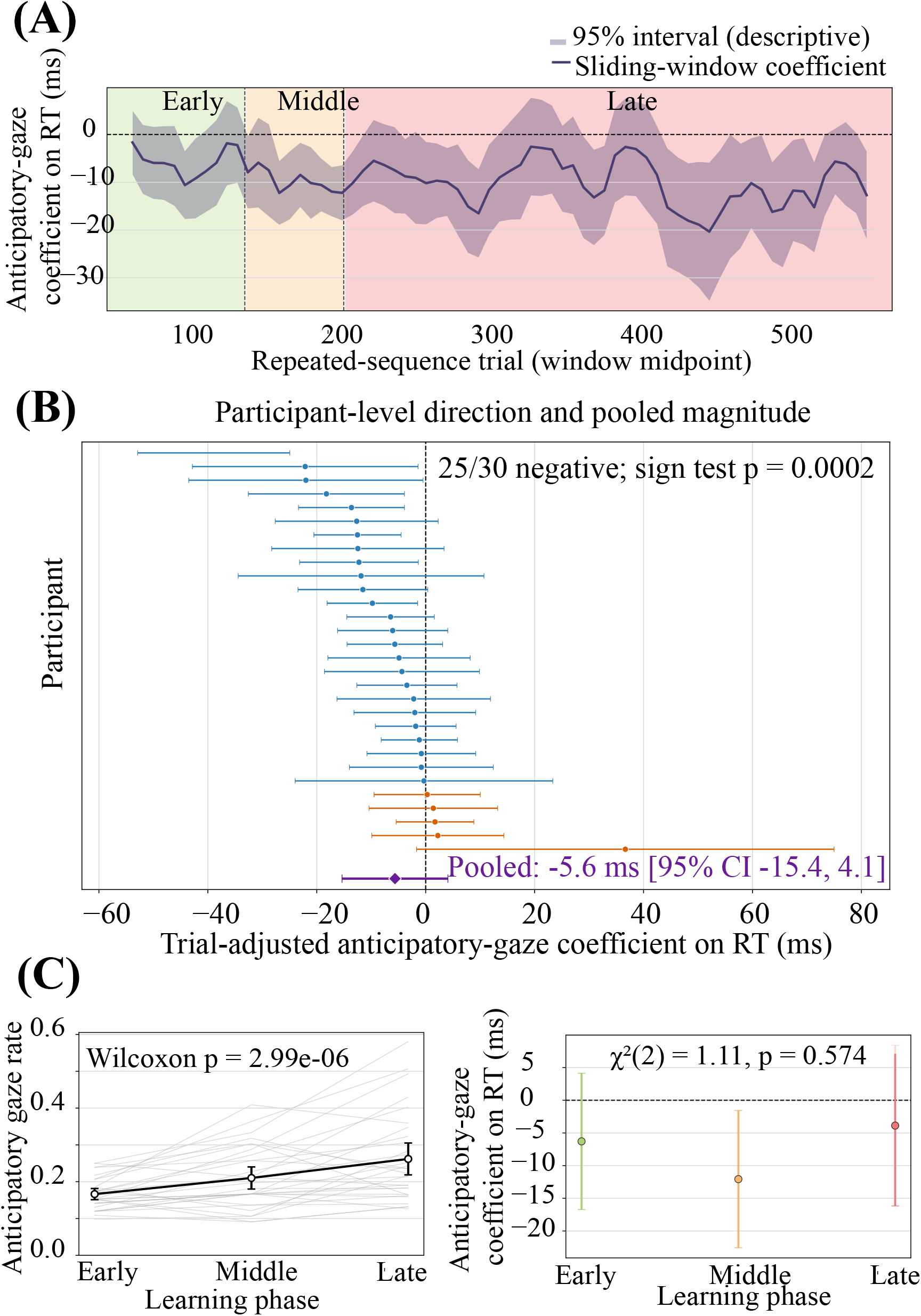
Association between anticipatory gaze and response time. (A) Sliding-window coeffiicients with S5% intervals, shown descriptively; inference derives from the trial-level model. (B) Participant-level trial- adjusted gaze coefficients with individual S5% intervals, sorted by magnitude, with the pooled participant- clustered estimate below. (C) Left: anticipatory gaze rate by phase, individual participants and group mean. Right: phase-specific gaze coefficients with S5% intervals from the trial-level model.

The average magnitude of this association however, was estimated with substantial uncertainty. Across participants, anticipatory-gaze trials were associated with responses that were 5.6 ms faster on average, but the confidence interval included zero (95% CI −15.4 to 4.1 ms; p = 0.256; Figure 4B). Thus, the participant-level estimates predominantly favoured faster responses on anticipatory trials, whereas the average magnitude of the association remained uncertain.

We found no evidence that the gaze–RT association changed across the three phases of the repeated-sequence period. Phase-specific estimates were −6.3 ms in the Early phase (95% CI −16.7 to 4.1 ms; p = 0.237), −12.1 ms in the Middle phase (95% CI −22.6 to −1.6 ms; p = 0.024), and −3.9 ms in the Late phase (95% CI −16.2 to 8.4 ms; p = 0.537). The joint gaze-by-phase interaction provided no evidence that the association differed across phases (χ²(2) = 1.11, p = 0.574; Figure 4C, right), and none of the pairwise phase contrasts was significant (Middle−Early = −5.8 ms, p = 0.412; Late−Middle = 8.2 ms, p = 0.297; Late−Early = 2.4 ms, p = 0.687). Although the Middle-phase estimate differed from zero when considered alone (p = 0.024), this does not indicate that the association differed between phases. We therefore interpret the association as not detectably different across phases, rather than as evidence that its magnitude remained constant across the repeated-sequence period. The sliding-window trajectory in Figure 4A is descriptive because consecutive windows overlap and was not used for statistical inference.

What did change across the repeated-sequence period was the frequency of anticipatory gaze. Mean anticipatory-gaze rate rose from 0.166 in the Early phase to 0.210 in the Middle phase and 0.262 in the Late phase, corresponding to a 57.5% increase from Early to Late. Twenty-four of 30 participants increased their anticipatory-gaze rate (Wilcoxon p = 2.99 × 10⁻⁶; Figure 4C, left). Thus, anticipatory gaze became progressively more frequent across the repeated-sequence period, whereas we found no detectable change in its trial-level association with response time.

### Exploratory phenotypes reveal distinct temporal organization of anticipatory gaze

We next explored whether participants could be grouped according to their patterns of anticipatory gaze and motor performance across the repeated-sequence period. Unsupervised clustering used six features describing each participant’s gaze and motor-performance profile: anticipatory-gaze rate in each of the three learning phases, improvement in RT from Early to Late, RT near the end of repeated-sequence practice, and skill score. This analysis identified three phenotypes: a higher-anticipation phenotype (n = 9), a typical phenotype (n = 18), and a small low-learning phenotype (n = 3; Figure 5A). A three-cluster solution provided the best separation of the candidate solutions tested (silhouette score = 0.353 for K = 3, compared with 0.320 for K = 2 and 0.339 for K = 4). The higher-anticipation phenotype showed the highest anticipatory-gaze rate averaged across the three learning phases, whereas the typical phenotype showed intermediate gaze rates and formed the largest group. The low-learning phenotype was characterized by poor sequence-specific performance together with RTs that worsened rather than improved across training (Figure 5B). Because these features were used to form the clusters, differences in those same features describe the phenotypes but do not provide independent evidence that the phenotypes differ.

**Figure 5.**
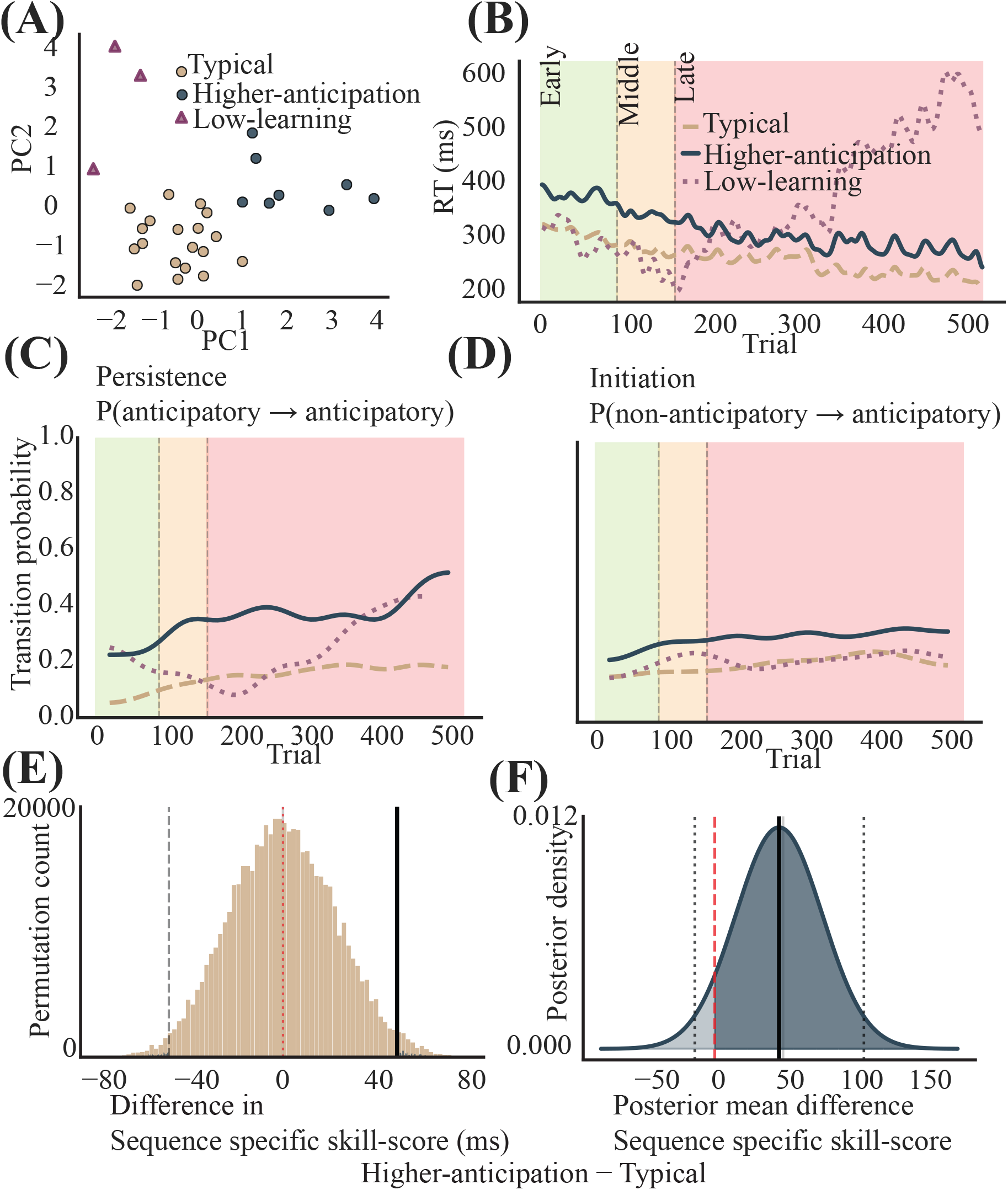
Learning phenotypes. (A) Principal component projection of the standardised six-feature space, coloured by phenotype. (B) Smoothed mean RT trajectory per phenotype across the repeated-sequence phase. (C) Persistence and (D) initiation transition probabilities per phenotype across practice.(E) Permutation null distribution and observed difference in sequence-specific skill score between the higher-anticipation and typical phenotypes. (F) Bayesian posterior for the same difference under a weakly informative prior.

We next examined whether the phenotypes differed in the trial-to-trial dynamics of anticipatory gaze. Averaged across the repeated-sequence phase, persistence—the probability that the next trial was also anticipatory given that the current trial was anticipatory—was 0.389 for the higher-anticipation phenotype compared with 0.152 for the typical phenotype, a difference of approximately 0.24. Initiation, the probability that the next trial was anticipatory given that the current trial was non-anticipatory, was 0.317 for the higher-anticipation phenotype versus 0.207 for the typical phenotype, a smaller difference of approximately 0.11 (Figure 5C–D). This difference was supported by the trial-level analysis: current gaze state predicted next-trial gaze differently in the higher-anticipation and typical phenotypes (OR = 1.83, p = 0.0012), indicating greater persistence in the higher-anticipation phenotype beyond its higher overall gaze frequency. Neither persistence nor initiation contributed to cluster formation, allowing these transition measures to characterize the resulting phenotypes using information not included in the clustering procedure.

The higher-anticipation phenotype also showed a 49.2-ms higher mean skill score than the typical phenotype (Figure 5E–F). The permutation and Bayesian summaries characterize the magnitude and uncertainty of this separation within the multidimensional phenotype solution. Because skill score was one of the six features used to construct the phenotypes, however, these comparisons are descriptive and should not be interpreted as independent evidence that phenotype membership is associated with greater sequence-specific learning.

Finally, we tested whether the persistence of anticipatory gaze, which distinguished the original phenotypes but was not used to form them, was itself associated with skill score. None of the three planned associations with skill score was reliable after correction for multiple comparisons: persistence across the session (ρ = 0.197, p = 0.306), Early-phase persistence (ρ = 0.194, p = 0.323), or initiation (ρ = 0.302, p = 0.111). Early-phase persistence also showed no independent association with skill score after accounting for participants’ Early-phase anticipatory-gaze rate (coefficient = −55.4 ms, permutation p = 0.689).

The two larger phenotypes therefore differed not only in how frequently anticipatory gaze occurred, but also in how consistently it was sustained across consecutive trials. This temporal organization provided information beyond the features used to define the phenotypes, although gaze persistence itself was not reliably associated with the magnitude of sequence-specific motor learning.

### Sensitivity analysis: excluding explicit learners

Because the study was designed to examine implicit sequence learning, we performed a sensitivity analysis to determine whether the principal findings were driven by participants who developed explicit knowledge of the repeating sequence. The five participants classified as explicit learners based on post-task awareness and correct reproduction of the full sequence were excluded (leaving *N* = 25), and the same preprocessing and analysis procedures were repeated in the reduced sample.

#### Behavioural learning

Evidence for sequence learning was preserved after explicit learners were excluded. Response time decreased from Random 1 to Early Repeat (*p* = 0.0045) and from Early Repeat to Late Repeat (*p* = 0.0254), before increasing when the repeated sequence was removed in Random 2 (*p* = 0.0022; all *n* = 25; Figure S1A). The mean skill score was 46.3 ms (SD = 67.6 ms), and 21 of 25 participants (84.0%) had positive scores, closely matching the proportion observed in the full sample (Figure S1B).

#### Between-participant divergence

Excluding the five participants who developed explicit knowledge did not change the main finding. Re-estimating the divergence points in the reduced sample showed that individual differences became detectable earlier in anticipatory gaze than in response time, with gaze diverging at Trial 116 and response time at Trial 158 (Figure S2A). The bootstrap analysis showed the same general ordering, with gaze diverging before response time in 90.3% of resamples (Figure S2B), although the estimated size of the temporal separation remained uncertain. Thus, the earlier differentiation of anticipatory gaze was not driven by participants who became explicitly aware of the sequence.

#### Learning phenotypes

The broad clustering pattern was also preserved after explicit learners were excluded. Repeating the clustering in the reduced sample again identified three phenotypes: a higher- anticipation phenotype (*n* = 6), a typical phenotype (*n* = 16), and a small low-learning phenotype (*n* = 3; Figure S3). Because skill score contributed to phenotype construction, differences in learning between these groups remain descriptive and were not interpreted as independent evidence that phenotype membership was associated with learning.

Together, these sensitivity analyses indicate that the behavioural evidence for sequence learning, the earlier differentiation of anticipatory gaze relative to response time, and the broad clustering pattern were qualitatively preserved after participants classified as explicit learners were excluded.

## Discussion

Implicit sequence learning is commonly inferred from changes in motor performance, although anticipatory gaze behaviour can provide information about the perceptual mechanisms involved in sequential motor learning^39,40^. In our study, between-participant differences became detectable earlier in anticipatory gaze than in response time. Across the repeated sequence, anticipatory gaze became progressively more frequent without a detectable change in its association with response time, and participants also differed in how consistently anticipatory gaze was sustained across consecutive trials. However, these gaze characteristics were not reliably associated with the magnitude of sequence-specific learning expressed at the end of the session. Together, the findings show that anticipatory gaze and motor performance provide related but non-redundant information about sequence learning: anticipatory gaze can provide a window into covert predictive processes that differentiate individuals before those differences become apparent in overt motor performance.

### Anticipatory gaze differentiates participants before motor performance

Individual differences became detectable in anticipatory gaze approximately 70 trials earlier than in response time, with gaze crossing the sustained divergence criterion at Trial 91 and response time at Trial 160. The bootstrap analysis favoured the same ordering in 95.5% of resamples. We therefore interpret this result as evidence for which measure revealed between-participant differences first, rather than as evidence for a fixed delay between perceptual and motor learning. The analysis does not suggest that perceptual learning began at Trial 91 or that motor learning began at Trial 160 but rather suggests that participants who remained comparatively similar in their motor performance had already begun to differ systematically in anticipatory gaze. Motor performance alone would therefore have missed an earlier source of individual variation in how upcoming elements of the sequence were anticipated.

Anticipatory eye movements provide a complementary measure of sequence learning because they reveal where participants expect an upcoming target before it appears. Previous serial reaction-time studies have shown that anticipatory eye movements reflect sequence learning and are associated with faster response times^26^,while implicit sequence-specific learning has also been demonstrated in purely oculomotor versions of the task without manual responding^41^. Anticipatory gaze can also reveal which elements of a sequence an individual has acquired and when those regularities become predictive^17^. More generally, trial-resolved modelling of sequence learning has shown substantial individual variation in how predictive sequence representations develop over practice^42^. More recently, Rubino et al. (2025)^8^ found that sequence-specific changes emerged earlier in oculomotor than in manual responses during implicit motor sequence learning. Complementary work in reaching has shown that motor sequence learning itself can dissociate improved prediction of what action comes next from optimization of how that action is executed^43^. Together, these findings suggest that gaze and response time provide related but non-redundant information about the acquisition and expression of sequential knowledge. Our findings extend this work from average changes in performance to the emergence of individual differences: anticipatory gaze distinguished participants while their response times remained comparatively similar.

### Anticipatory gaze becomes more frequent across trials without a detectable change in its association with response time

At the trial level, anticipatory gaze and motor performance were related, but the magnitude of this relationship was small and uncertain. Twenty-five of 30 participants responded faster on trials in which gaze was already at the upcoming target, indicating a consistent direction of association across individuals. However, the average difference was small (5.6 ms) and its confidence interval included zero. Anticipatory gaze was therefore commonly associated with faster responses within individuals, but the magnitude of this relationship was uncertain. Trials containing anticipatory gaze therefore also tended to contain faster motor responses, but this association does not establish that the eye movement itself facilitated the response. Both may instead reflect a common predictive process through which information about the upcoming spatial goal becomes available before target onset^44^.

What changed more clearly across trials was the frequency with which predictive gaze was expressed. Anticipatory gaze increased by 57.5% from the Early to Late phase, with increases observed in 24 of 30 participants. In contrast, we found no evidence that the gaze–response time association differed across phases. This pattern suggests that progression across repeated-sequence trials was characterized primarily by more frequent expression of advance prediction, rather than by a detectable increase in the response time advantage associated with each anticipatory event. . Importantly, the absence of a gaze-by-phase interaction does not establish that the relationship remained constant; rather, any change was not detectable in the present data.

The increasing frequency of anticipatory gaze may reflect predictive information being expressed more consistently as participants progressed across repeated-sequence trials. Similar changes toward predictive gaze have been observed during visuomotor skill acquisition, where eye movements shift from tracking ongoing actions toward upcoming spatial goals^16,45^. In sequence-learning tasks, anticipatory eye movements can reveal where participants expect the next target and aspects of predictive knowledge that are not apparent from response time alone.Oculomotor anticipation may also become more likely across sequence exposure independently of learning which specific target comes next^10^. A similar dissociation has been observed in spatial sequence memory, where anticipatory eye movements increased across repetitions but their expression did not directly track sequence-learning magnitude^27^. Thus, greater anticipatory-gaze frequency need not produce an increasingly large response time advantage: progression across repeated-sequence trials may increase how consistently predictive information is expressed without proportionally changing its relationship with the subsequent motor response. Whether this reflects greater availability of predictive information or a greater tendency to deploy it cannot be distinguished from the present data.

### Individual differences extend to the temporal organization of anticipatory gaze

The individual differences revealed by anticipatory gaze extended beyond how often participants anticipated upcoming targets to how that behaviour unfolded across consecutive trials. Among the two larger exploratory phenotypes, higher-anticipation participants were more likely than typical participants to remain anticipatory after an anticipatory trial. This distinction in persistence was not used to define the phenotypes, suggesting that differences in overall anticipatory-gaze frequency were accompanied by differences in the temporal organization of predictive behaviour. Rather than differing only in whether prediction occurred, participants therefore also differed in how consistently anticipatory behaviour was sustained from one trial to the next.

This temporal organization may reveal information that is obscured by average gaze frequency alone. Sequence learning is increasingly understood as heterogeneous: individuals can acquire different subsets of sequential regularities, and similar overall performance can arise from different underlying learning processes^46,47^ . Our findings extend this heterogeneity to the continuity of predictive behaviour. Two participants with similar overall levels of anticipation could differ in whether predictive gaze occurs sporadically or is sustained across consecutive elements of the sequence. Persistence may therefore provide a complementary description of how predictive information is expressed over time, although the present data do not establish the cognitive mechanism underlying that stability.

Importantly, these differences in the organization of predictive gaze were not reliably associated with how much sequence-specific motor learning participants ultimately expressed. Although the original phenotypes differed descriptively in skill score, that outcome contributed to their definition and therefore could not provide independent evidence of a phenotype–learning relationship. Persistence and initiation likewise showed no reliable associations with skill score. Predictive gaze and motor learning should therefore not be treated as interchangeable expressions of a single learning process: participants may differ substantially in how they anticipate upcoming events without those differences mapping directly onto the magnitude of their motor learning. This interpretation is consistent with evidence that oculomotor and manual expressions of sequence learning can dissociate. Sequence-specific learning can be expressed in a purely oculomotor task without a manual response^41^, and response time and correct oculomotor anticipations can show different patterns of offline change^48^. More recently, sequence-specific learning effects in saccades and reaches were not correlated at retention despite learning being evident in both systems^8^. Thus, weak sequence-specific motor performance does not necessarily imply an absence of predictive information; perceptual prediction and its expression in motor performance may vary partly independently.

### Limitations

Several limitations constrain the interpretation of these findings. The Early, Middle, and Late phase boundaries were derived from the same dataset in which subsequent phase-based analyses were performed; independent replication will therefore be important for establishing the timing of gaze and response time differentiation. As well, the clustering analysis was exploratory and the smallest phenotype contained only three participants, so the resulting phenotype structure should not be interpreted as evidence for discrete learner types^49^. Finally, the divergence points describe when between-participant heterogeneity became detectable at the group level, not when perceptual or motor learning began within individual participants. Trial 91 and Trial 160 should therefore be interpreted as population-level behavioural landmarks rather than individual learning thresholds.

### Future Directions

The interval between gaze and RT differentiation provides a testable window for identifying the processes that precede overt motor differentiation. Sequence performance can improve through changes in planning both before and during movement execution, rather than through motor execution alone^50^.. Neural changes during sequential motor learning can also be tracked relative to individual behavioural markers of emerging sequence knowledge, revealing person-specific temporal changes in frontoparietal network involvement^51^. Future work combining eye tracking with temporally resolved neural measures could therefore test whether neural signatures of sequence prediction and action preparation differentiate during the period in which anticipatory gaze already distinguishes participants but response time does not. This would help determine whether the behavioural ordering observed here reflects a corresponding progression in the processes supporting prediction, action planning, and motor expression. As well, participants encountered a single repeated sequence within one session. Future studies can investigate whether the temporal ordering seen here generalizes across sequence structures or persists across consolidation and longer-term learning^52,53^.

### Conclusion

In conclusion, anticipatory gaze revealed individual differences in implicit sequence learning before those differences became apparent in response time. Anticipatory gaze became increasingly frequent across the repeated sequence and differed between participants in its persistence across consecutive trials, although these gaze characteristics were not reliably associated with the magnitude of sequence-specific motor learning. Sequence learning is therefore not fully captured by a single motor-performance trajectory: predictive behaviour and motor expression can differentiate participants at different times and in different ways. Concurrent measurement of both perceptual and motor domains may provide a more complete account of how sequential knowledge emerges during the learning process and is expressed through action.

## Supplemental Figures

**Figure S1:**
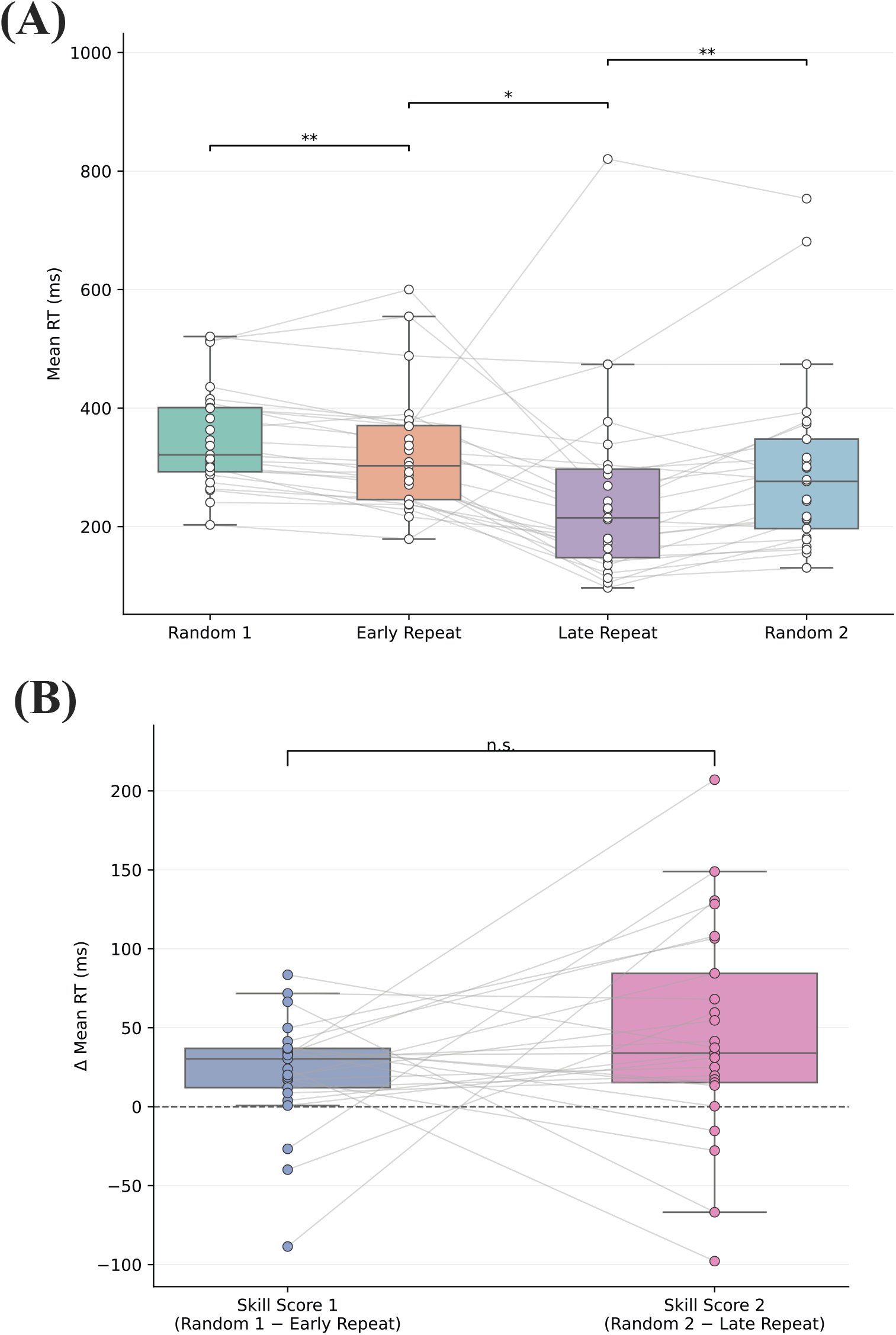
Behavioural learning after excluding explicit learners. (A) Mean response time across Random 1, Early Repeat, Late Repeat, and Random 2 in the reduced sample (N = 25), with individual participant trajectories. (B) Early and late sequence-specific skill scores for individual participants. Positive scores indicate faster responses during the repeated sequence relative to the corresponding random block.

**Figure S2:**
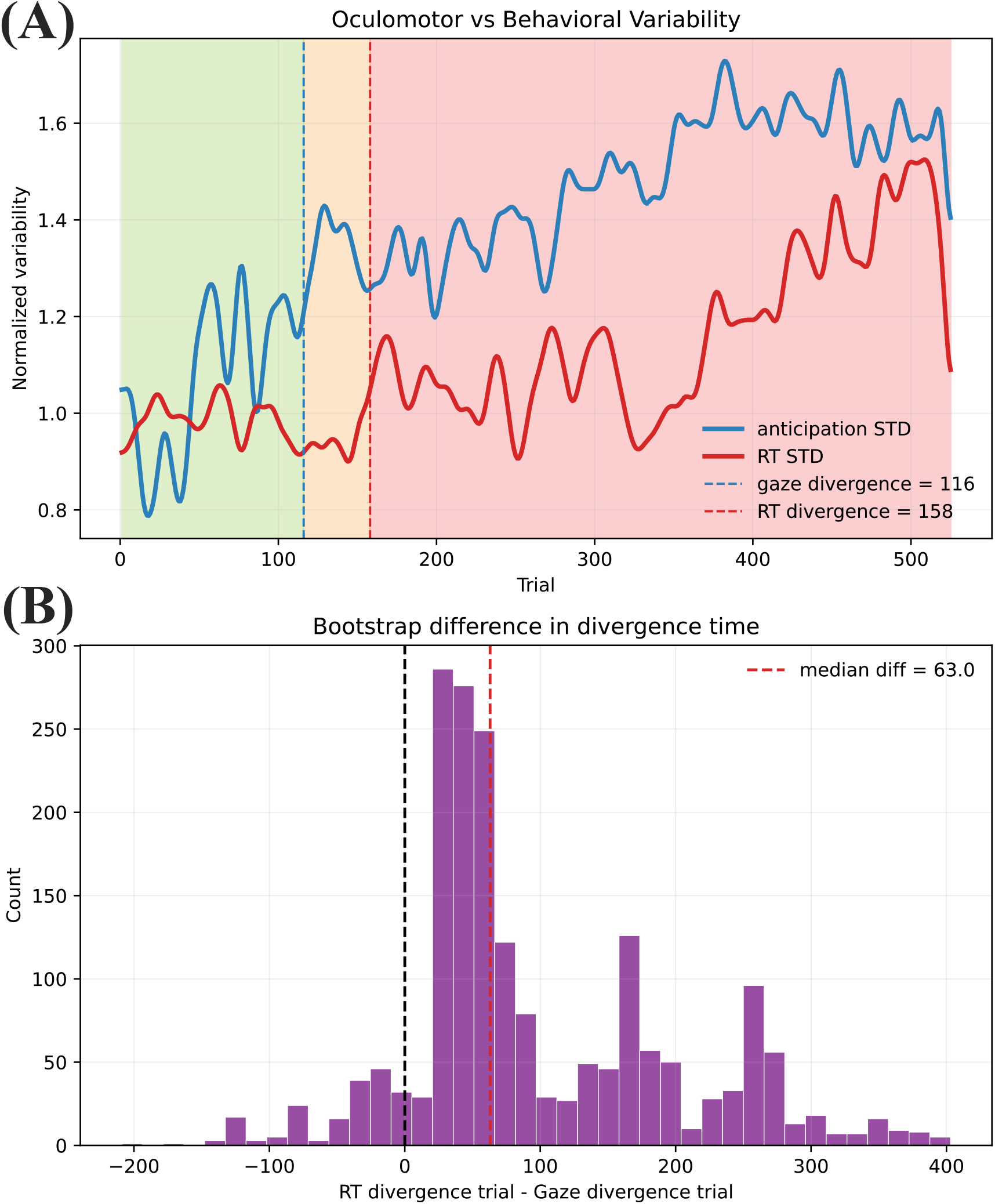
Between-participant divergence after excluding explicit learners. (A) Normalised between-participant variability in anticipatory gaze and response time across repeated-sequence trials. Anticipatory gaze diverged at Trial 116 and response time at Trial 158. (B) Bootstrap distribution of the difference between response-time and gaze divergence points; positive values indicate earlier divergence of anticipatory gaze.

**Figure S3:**
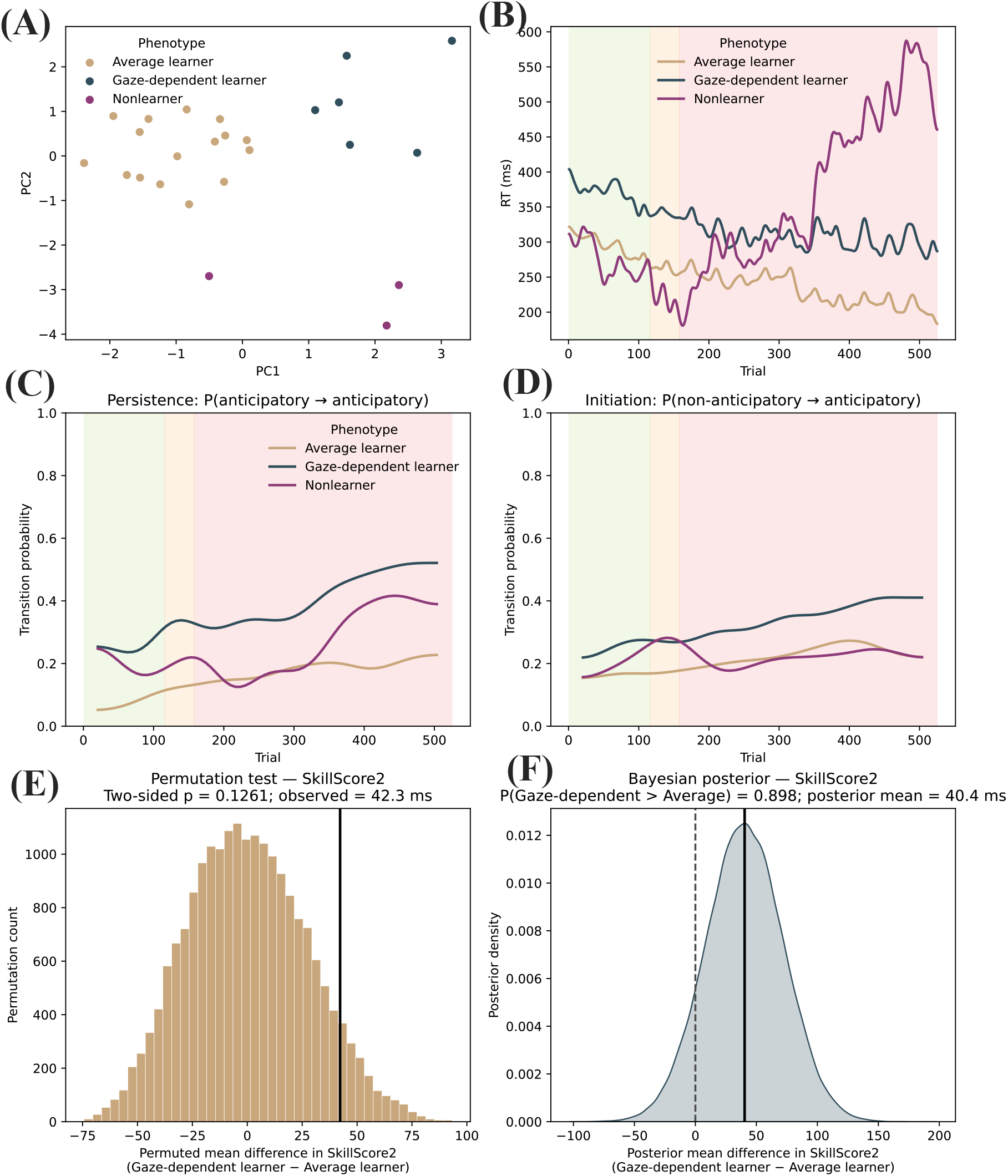
Learning phenotypes after excluding explicit learners. (A) Principal-component projection of the clustering feature space. (B) Mean response-time trajectories across repeated-sequence trials by phenotype. (C) Persistence and (D) initiation probabilities across trials. (E) Permutation distribution of the between-phenotype difference in sequence-specific skill score. (F) Bayesian posterior distribution of the same difference.

