## Supplementary figures and images for "Anticipatory Gaze Reveals Individual Differences in Implicit Sequence Learning Before Motor Performance"

### Supplemental figure 1

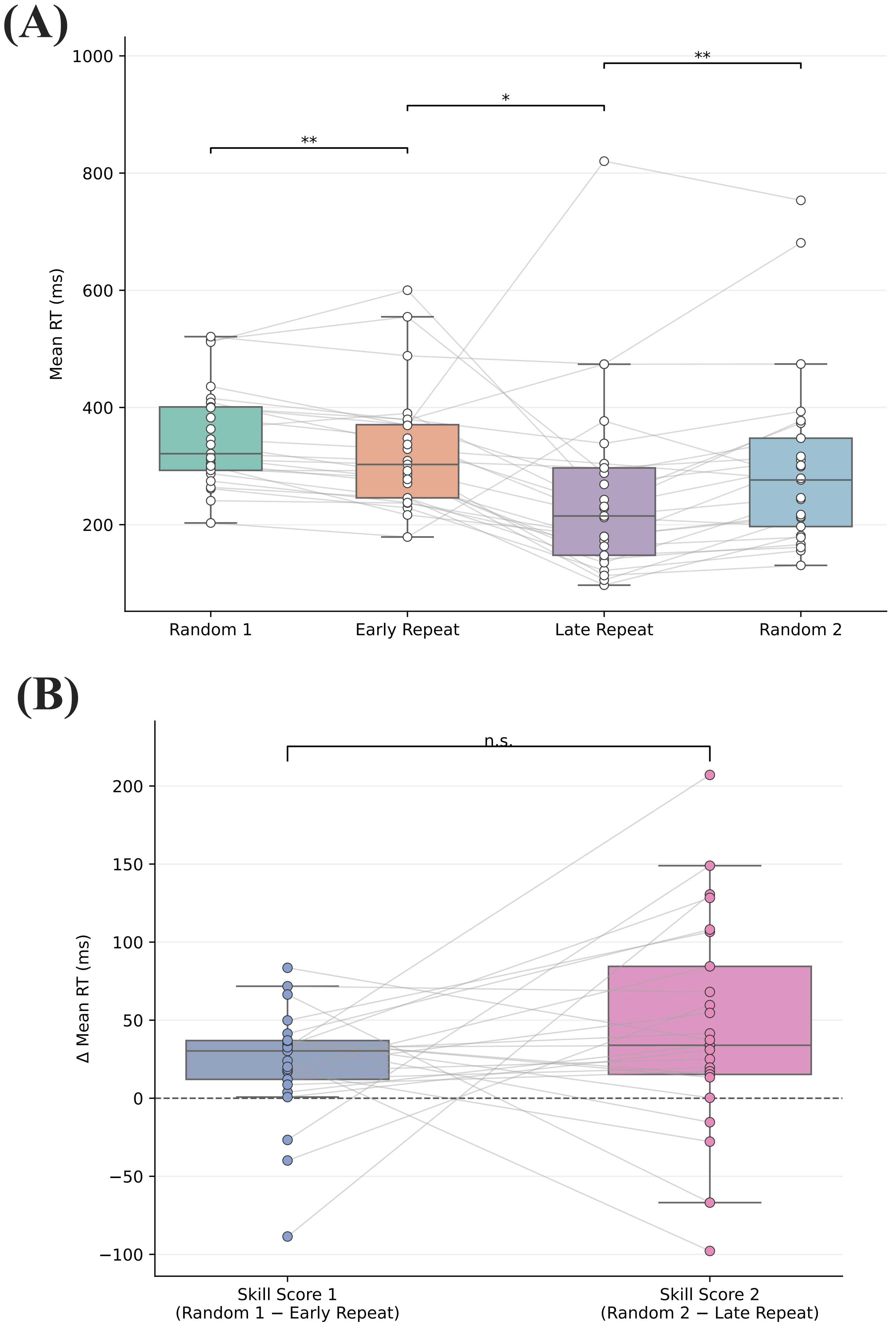

### Supplemental figure 2

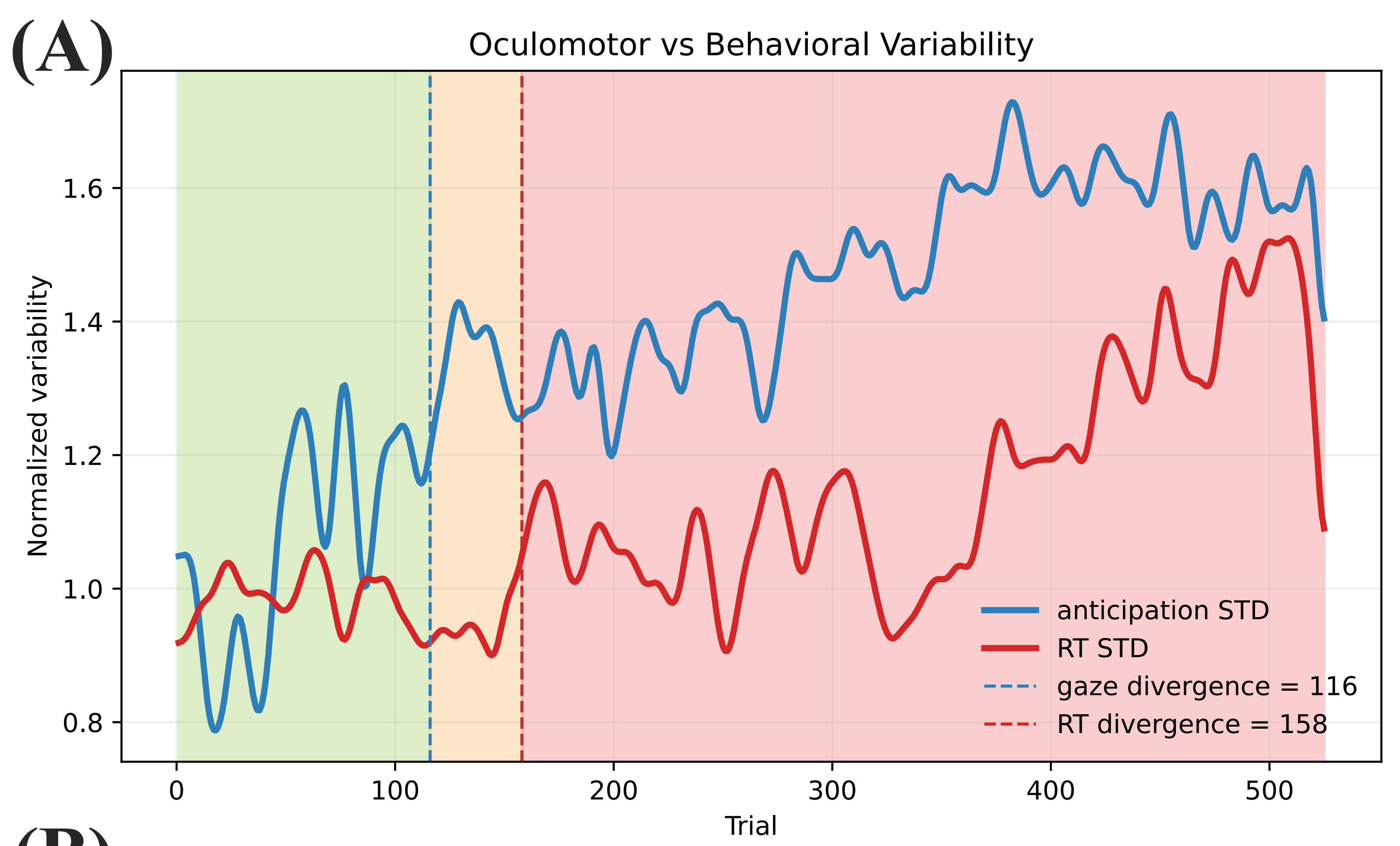

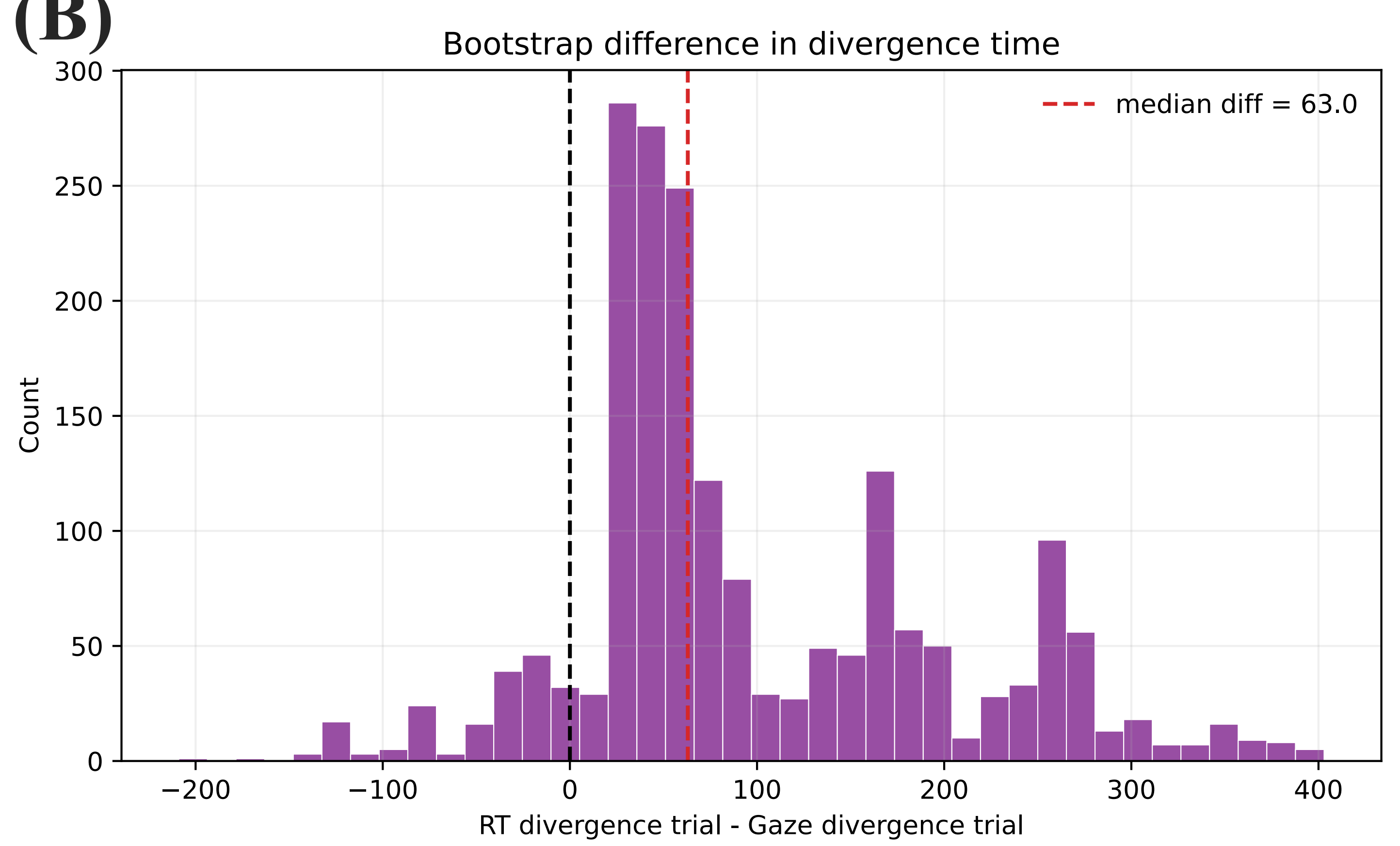

### Supplemental figure 3

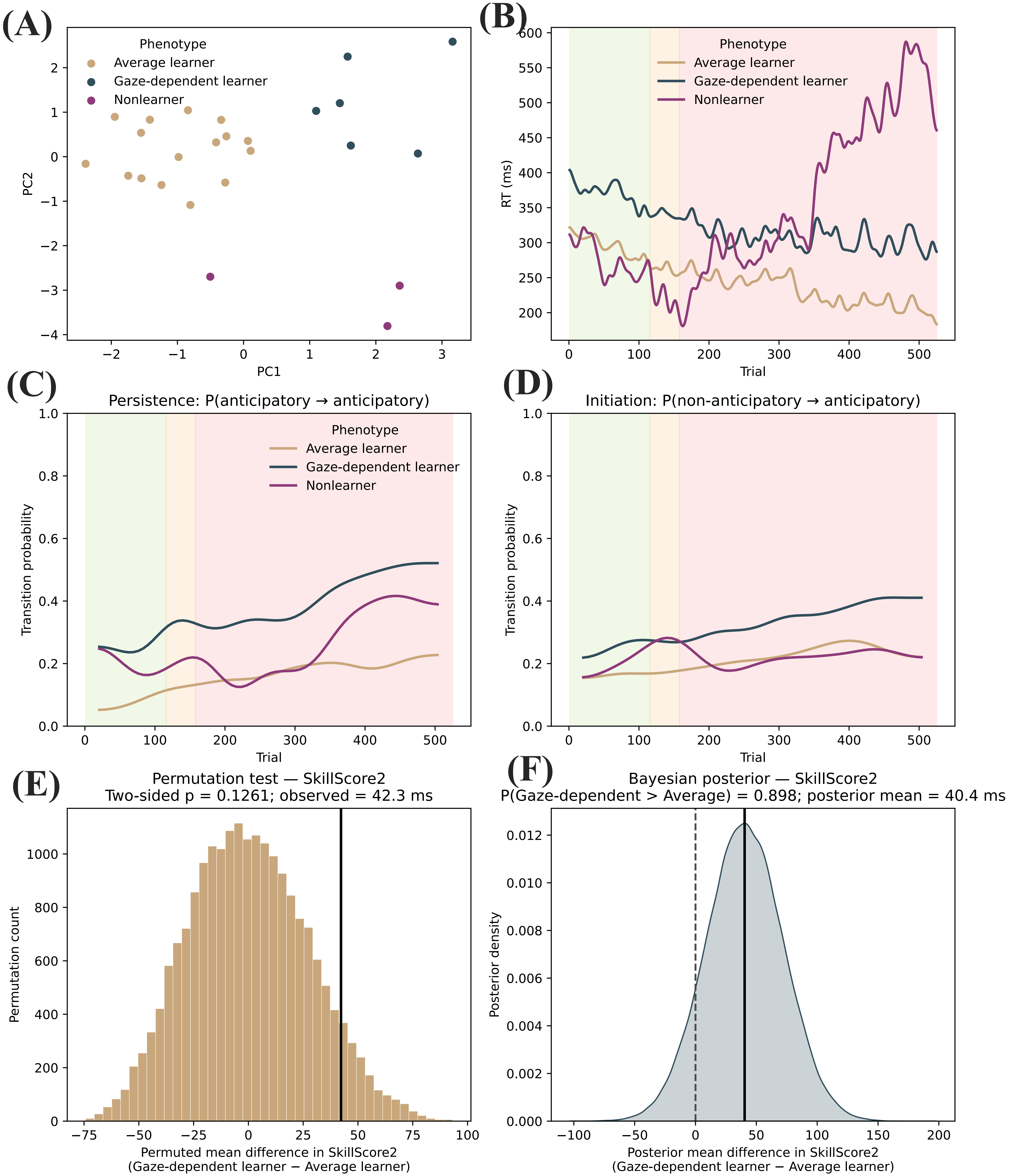
